# Systematic identification of exercise-induced anti-aging processes involving intron retention

**DOI:** 10.1101/2024.04.25.591048

**Authors:** Hayata Kodama, Hirotaka Ijima, Yusuke Matsui

## Abstract

Exercise is one of the most promising anti-aging interventions for maintaining skeletal muscle health in older adults. Nine “Aging Hallmarks”, proposed by López-Otín, offer insights into the aging process; however, the link between these hallmarks and exercise is not fully elucidated. In this study, we conducted a systematic multi-omics analysis of skeletal muscles, focusing on aging and exercise, based on gene signatures for aging hallmarks. It is posited that mRNA splicing activity, linked to genomic instability, constitutes a fundamental hallmark of aging, and it exhibits divergent expression patterns in response to aging and exercise. Additionally, we analysed splicing events and discovered that intron retention (IR) is significantly impacted by aging, exhibiting contrasting changes to those induced by resistance training in the older cohort. The isoforms characterised by IR are notably enriched in mitochondrial functions. Conclusively, our results underscore the significance of splicing mechanisms as a novel aspect of aging hallmarks in skeletal muscles and propose a new mechanism by which exercise exerts its anti-aging effects on skeletal muscles through intron retention.

**Key points summary:**

- Skeletal muscle aging involves significant structural and functional changes, including loss of muscle mass, decline in strength, and mitochondrial dysfunction, all influenced by genomic instability.
- Exercise has been identified as a key intervention that counters genomic instability and modulates mRNA splicing patterns, particularly through the regulation of Intron Retention, to mitigate aging effects in skeletal muscle.
- We reveal the novel role of IR, especially in principal isoforms, where it is linked to critical cellular processes like mitochondrial function, suggesting a targeted pathway through which exercise exerts its anti-aging effects.
- The findings provide new insights into the molecular mechanisms underlying the beneficial effects of exercise on aging skeletal muscle.
- This study lays the groundwork for future research on exercise-induced modulation of mRNA splicing as a therapeutic strategy for aging and potentially age-related diseases, pointing towards a significant shift in how we approach aging intervention strategies.

## Introduction

Skeletal muscle aging is primarily characterised by a loss of muscle mass and a decline in muscle strength (Morley *et al*., 2001). Additionally, it leads to various structural and functional changes, including a reduction in type II muscle fibre size (Nilwik *et al*., 2013) and motor units (Kung *et al*., 2014; Piasecki *et al*., 2016), anabolic resistance (Cuthbertson *et al*., 2005), mitochondrial dysfunction (Short *et al*., 2005; Chabi *et al*., 2008), and a decline in muscle regenerative capacity (Shefer *et al*., 2010; Chakkalakal *et al*., 2012). Among the myriad of promising anti-aging interventions for improving skeletal muscle health, exercise elicits systemic effects, primarily through muscle contraction, and ameliorates various functional impairments associated with skeletal muscle aging, including reductions in muscle mass (Peterson *et al*., 2011), strength (Lu *et al*., 2021), mitochondrial function (Lanza *et al*., 2008), and muscle regeneration potential (Shefer *et al*., 2010). However, the detailed molecular mechanisms underpinning these anti-aging effects of exercise are not fully elucidated, presenting a significant challenge in optimising exercise-based intervention strategies considering aspects such as modality, intensity, and duration.

Recent proteomic analyses of healthy human skeletal muscles have indicated that some age-related changes in mitochondrial and mRNA splicing-related protein levels (Ubaida-Mohien *et al*., 2019*b*) are reversed in master athletes (Ubaida-Mohien *et al*., 2022) and individuals with high physical activity (Ubaida-Mohien *et al*., 2019*a*). Additionally, meta-analyses of an extensive collection of public transcriptome data on human skeletal muscle have revealed pathways and upstream regulators induced by exercise, a subset of physical activity (Pillon *et al*., 2020; Amar *et al*., 2021). Nonetheless, the precise molecular-level impacts of exercise on healthy aging necessitate further investigation. An integrated analysis of multi-omics data concerning aging and exercise could provide novel insights into the molecular mechanisms driving the anti-aging effects of exercise on skeletal muscles.

One pivotal clue in deciphering the anti-aging effects of exercise on skeletal muscles lies in the concept of aging hallmarks (López-Otín *et al*., 2013). Aging hallmarks represent the fundamental aspects of aging across nine biological dimensions: genomic instability, telomere attrition, epigenetic alterations, loss of proteostasis, deregulated nutrient sensing, mitochondrial dysfunction, cellular senescence, stem cell exhaustion, and altered intercellular communication (López-Otín *et al*., 2013) (**Figure 1A**). Previous research has established connections between aging hallmarks and the typical clinical manifestations of aging (Aunan *et al*., 2016), the co-occurrence of age-related diseases (Fraser *et al*., 2022), and skeletal muscle-specific aging (Wiedmer *et al*., 2021). Notably, exercise is posited to influence all aging hallmarks (Garatachea *et al*., 2015).

**Figure 1.**
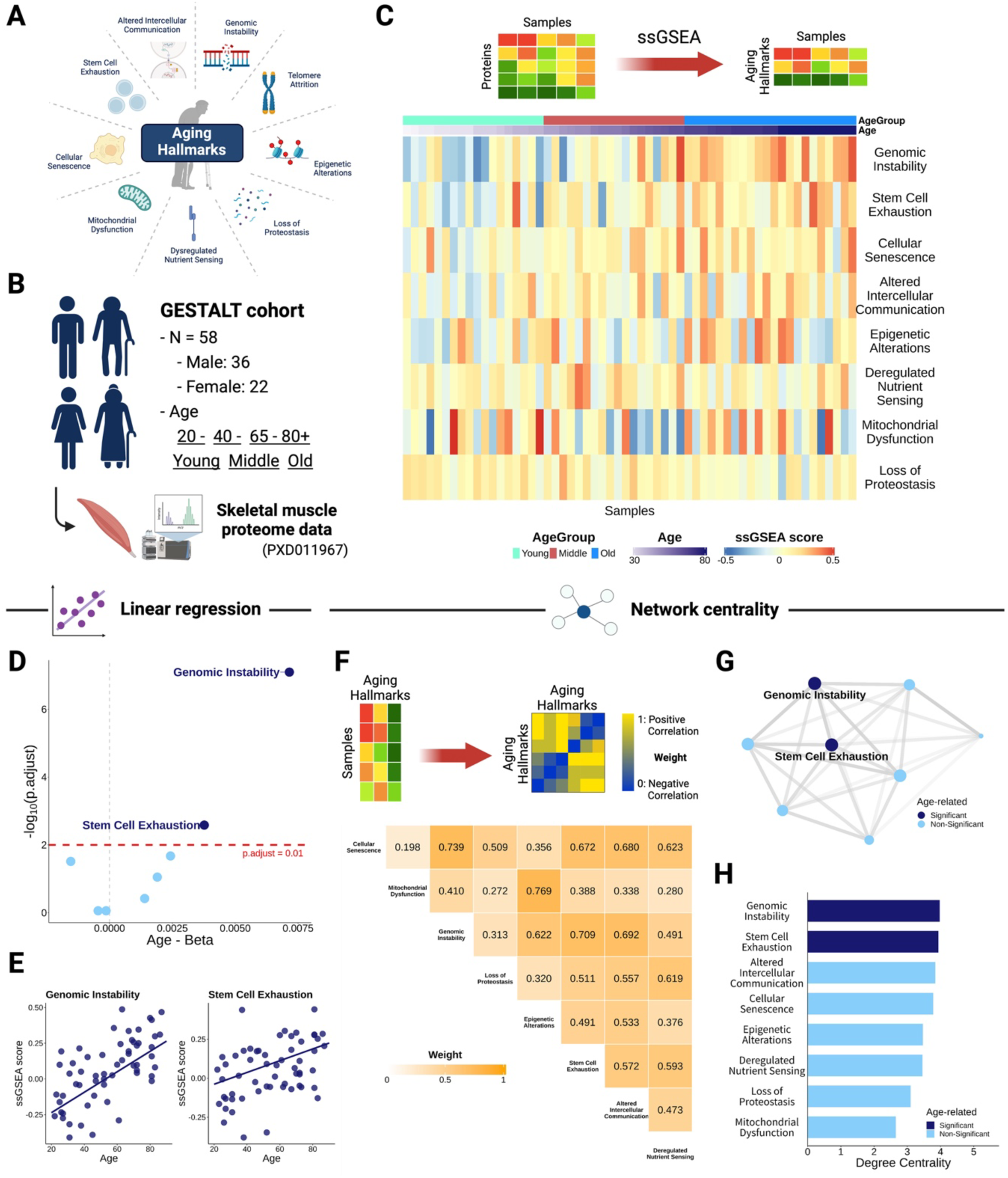
Genomic instability as a leading hallmark of skeletal muscle aging. **A)** Overview of nine aging hallmark categories. **B)** Details of the GESTALT cohort dataset, which includes a range of ages, used for analysing healthy aging. **C)** A heatmap displaying the activities of aging hallmarks as estimated using ssGSEA, with enrichment scores based on each hallmark’s molecular signature. Rows represent aging hallmarks, and columns represent samples, organised by age from youngest to oldest. **D)** A volcano plot illustrating the relationship between aging hallmarks and skeletal muscle aging from linear regression analysis. The x-axis shows the aging dependency of each hallmark as indicated by the regression coefficient for age, while the y-axis shows the degree of statistical significance, represented by the negative log10 of the adjusted p-value for the regression coefficient. The red dashed line denotes a 1% significance threshold. **E)** The relationship between age and the activities of genomic instability and stem cell exhaustion hallmarks, with age on the x-axis and hallmark enrichment on the y-axis, adjusted for sex. The plotted lines represent regression lines. **F)** A heatmap of correlations among aging hallmarks, with correlation strengths derived from the enrichment magnitudes estimated using ssGSEA and rendered as unsigned values. **G)** A weighted network diagram illustrating the relationships between aging hallmarks based on correlation coefficients from F. H) A representation of the interconnectedness among aging hallmarks using a weighted network centrality measure to indicate each hallmark’s significance, highlighted in dark blue for those significantly associated with skeletal muscle aging as shown in E. Parts of the figures were generated with biorender.com.

In this study, we conducted an integrated analysis of transcriptomic and proteomic data on skeletal muscle aging and exercise, guided by an aging hallmark-focused systematic approach. Our findings reveal that genomic instability, closely associated with mRNA splicing, stands as the principal hallmark of skeletal muscle aging. Moreover, we observed that exercise attenuates mRNA splicing activities. Through splicing event analysis, we identified that intron retention (IR) significantly altered with aging and was inversely related to the effects of resistance training (RT) in the older cohort. Enrichment analysis of IRs impacting isoform expression highlighted mitochondrial functions, especially within the electron transport chain. The findings of this study imply that modifications in splicing regulation, particularly those induced by IR, are crucial for the anti-aging benefits of exercise.

## Materials and methods

### Retrieval of preprocessed human skeletal muscle proteome data

We compiled preprocessed human skeletal muscle proteome data (PXD011967) of a very healthy cohort of individuals across a wide age range (20 to 80 years) from the Genetic and Epigenetic Study of Aging and Laboratory Testing (GESTALT) (Ubaida-Mohien *et al*., 2019*b*). The age distribution of subjects was 20-34 (n=13), 35-49 (n=11), 50-64 (n=12), 65-79 (n=12), and 80+ (n=10) years of age, taken from the lateral vastus muscle. We performed missing-value filtering, wherein proteins were quantified using at least five samples from each age group. The three age groups were defined as follows: young (< 40 years), middle (40–65 years), and old (> 65 years). We then imputed missing values using the R package missForest (Stekhoven & Bühlmann, 2012). Finally, 4,289 proteins were used for downstream analysis.

### Characterisation of skeletal muscle aging based on aging hallmarks

To characterise skeletal muscle aging, we evaluated the aging hallmarks using two approaches. Initially, we compiled gene signatures representing each hallmark from the Aging Atlas database (Aging Atlas Consortium, 2021), focusing on hallmarks with a minimum of three quantified proteins and excluding telomere attrition. Next, a single-sample Gene Set Enrichment Analysis (ssGSEA) (Barbie *et al*., 2009) was performed using the R/Bioconductor package GSVA (Hänzelmann *et al*., 2013) to estimate the activity of specific gene sets in individual samples and assess their association with skeletal muscle aging. Linear regression models were used to analyse the relationship between each hallmark and skeletal muscle aging, with hallmark activity estimated via ssGSEA as the dependent variable, and age and sex as independent variables.

Furthermore, a network analysis was conducted to identify central hallmarks influencing other aspects of skeletal muscle aging. Utilising hallmark activities estimated via ssGSEA, a network was constructed, and network centrality was assessed. Edges in the network were weighted based on Pearson’s correlation, adjusting the scale from [-1, 1] to [0, 1] to maintain the sign of each correlation, where 0 indicates a negative correlation, and 1 indicates a positive correlation. We selected the degree centrality, which is the sum of edge weights for each hallmark, to measure network centrality.

### Exploration of the pathways affected by genomic instability using directional GSEA

Directional GSEA (dGSEA) is an extension of GSEA that leverages gene ranks based on their association with a specific gene set (source) to evaluate the influence of other gene sets (targets) comprehensively. The weights of the gene ranks are the sum of Pearson’s correlation with the source in each target. In this analysis, 15 proteins related to genomic instability served as the source, and 4,274 other proteins functioned as targets. Target gene ranks were analysed using standard GSEA algorithms implemented in the R/Bioconductor package ClusterProfiler. Pathway-related gene sets were derived from the Reactome Pathway database (Gillespie *et al*., 2022) for pathway analyses. Additionally, to explore the association between mRNA splicing, as influenced by genomic instability, and aging, we determined eigenproteins through Principal Component Analysis (PCA) of the expression data for 78 proteins in the leading edge of mRNA splicing. Associations with age were evaluated using Pearson’s correlation.

### Human skeletal muscle transcriptome data pre- and post-exercise training and gene expression quantification

We compiled human skeletal muscle RNA-seq data (GSE97084) of vastus lateralis derived from a healthy cohort, both pre- and post-exercise training, including young (18-30 years old; n=29) and older (65-80 years old; n=23) participants (Robinson *et al*., 2017). All participants undertook one of the following exercise modalities for 12 weeks: high-intensity interval training (HIIT), RT, or combined training (CT). One post-exercise sample was missing for 52 participants. Other sample details have been provided in a previous study. Raw data were obtained from the Sequence Read Archive (SRA) in FASTQ format. Gene expression was quantified using the nf-core/rnaseq pipeline, which aligned sequencing reads to the hg38 reference genome using STAR (Dobin *et al*., 2013) and quantified gene expression using Salmon (Patro *et al*., 2017).

### Differential gene expression analysis and GSEA

The differential gene expression analysis pre- and post-exercise training was conducted according to the workflow described by Law et al (Law *et al*., 2016). This involved filtering out genes with low expression, applying trimmed mean of M-values (TMM) normalisation (Robinson & Oshlack, 2010), and using Voom transformation (Law *et al*., 2014) as preprocessing steps before comparing pre- and post-exercise training gene expression levels. Moreover, six subgroup analyses were performed, with subgroups defined based on two age group categories –young and older– and three exercise modalities: HIIT, RT, and CT. Gene ranks derived from t-statistics in the differential gene expression analysis were input into GSEA, using gene sets related to pathways. The R/Bioconductor packages edgeR (Robinson *et al*., 2010), limma (Ritchie *et al*., 2015), and clusterProfiler (Wu *et al*., 2021) were employed for these analyses.

### mRNA splicing-related factors based on the leading edge of GSEA

Genes within the leading edge of GSEA contribute significantly to the enrichment of a gene set and are considered to be associated with the GSEA target. Similarly, genes within the leading edge of dGSEA are linked to the source, in this case, proteins related to genomic instability. This study focused on investigating mRNA splicing-related factors based on the leading edges of GSEA and dGSEA. We analysed overlapping mRNA splicing-related proteins that are upregulated in conjunction with genomic instability during skeletal muscle aging and mRNA splicing-related genes that are downregulated following exercise. Additionally, we assessed the bias of these overlapping factors towards spliceosomes, which are mainly responsible for mRNA splicing. Gene sets pertaining to spliceosomes were sourced from the Gene Ontology project (Ashburner *et al*., 2000; Gene Ontology Consortium *et al*., 2023).

### Splicing event analysis focusing on IR

We analysed all splicing events, including skipping exons (SE), mutually exclusive exons (MX), alternative 5’/3’ splice sites (A5/A3), alternative first/last exons (AF/AL), and IR, using human skeletal muscle RNA-seq data (GSE129643) collected from a very healthy cohort in GESTALT (Ubaida-Mohien *et al*., 2019*b*). Percent Spliced In (PSI) values were calculated using SUPPA2 (Trincado *et al*., 2018), with isoform-level transcripts per million (TPM) values generated by the nf-core/rnaseq pipeline (Ewels *et al*., 2020) as input. PSI values quantify the proportion of isoforms undergoing a specific splicing event relative to all isoforms for a given gene. Splicing events were identified using GTF files from the Ensembl hg38 v110 database. Datasets devoid of missing values were used in downstream analyses. We assessed the relationship between each splicing event and healthy aging using linear regression, with PSI values as the dependent variable, and age and sex as independent variables.

Focusing on IR, we also explored splicing modifications related to aging and exercise by analysing human skeletal muscle RNA-seq data (GSE97084) collected pre- and post-exercise training (Robinson *et al*., 2017). We examined age-related increases in IR by comparing pre-exercise training samples from young and older groups. Mean PSI values between groups, both pre- and post-exercise, and across six subgroups delineated by age and exercise modality were compared to assess IR changes.

### IR categorisation based on principal isoforms

To further investigate IR alterations due to aging and exercise, we classified IR into three categories: IR-P (included in principal isoforms), IR-A (not included in principal isoforms), and others. Principal isoforms, labeled ‘PRINCIPAL: 1’ through ‘PRINCIPAL: 5’ in the APPRIS database (Rodriguez *et al*., 2022), were selected for analysis. Given that multiple principal isoforms may share the same coding sequence (CDS) for a single gene in the APPRIS database and are likely functionally similar at the protein level, all such isoforms were included in our analysis. We investigated age- and exercise-related changes in IR, IR-P, and IR-A through linear regression analysis or by comparing mean PSI values between the groups, such as pre- and post-exercise training.

### Analysis of the association between IR and principal isoform expression

To ascertain the relationship between IR and changes in isoform expression, we analysed Pearson’s correlation between changes in IR-P or IR-A and principal isoform expression due to aging and exercise. In cases where multiple principal isoforms corresponded to an IR, the isoform with the largest absolute change in biological effect was chosen for analysis. Changes in principal isoform expression were quantified using TPM value shifts, i.e., the regression coefficient for age from the linear regression analysis, adjusted for sex, or the log2 fold change (logFC) between groups.

### Enrichment analysis of ‘influential IR’

We aimed to identify biological functions associated with IR by detecting ‘influential IR’, whose changes with aging and exercise significantly impact principal isoform expression. For aging, within each cohort, a decrease in IR-P or an increase in IR-A with downregulated principal isoforms were tagged as ‘influential IRs’. Conversely, for exercise (specifically RT in the older group), ‘influential IRs’ were defined by an increase in IR-P or a decrease in IR-A with upregulated principal isoforms. These overlapping IRs not only exhibit divergent changes during aging and exercise but also significantly influence principal isoform expression. Enrichment analysis was conducted using the R/Bioconductor fedup package.

### Statistical analysis

All statistical analyses were executed using R version 4.1.2. Either linear regression or Fisher’s exact test was employed for statistical analyses. The Benjamini–Hochberg method was used to adjust p-values for multiple comparisons, with adjusted p-values < 0.01 deemed statistically significant.

## Results

### Genomic instability signatures correlate significantly with skeletal muscle aging

We investigated the age-dependent aging hallmarks in skeletal muscle. To minimise confounding factors unrelated to aging, we selected a very healthy aging cohort. We used skeletal muscle proteome data of healthy adults across their lifespan (22–87 years old; n = 58) from the GESTALT cohort study (PXD011967) (Ubaida-Mohien *et al*., 2019*b*) (**Figure 1B**). Additionally, we compiled the molecular signatures characteristic of each hallmark from the Aging Atlas database (Aging Atlas Consortium, 2021). The activity level of each hallmark within an individual was estimated using ssGSEA (Barbie *et al*., 2009), which determines the enrichment score of a molecular signature based on the expression level in each sample (**Figure 1C**). We then conducted a regression analysis, incorporating age and sex as explanatory variables for each estimated hallmark activity, to identify hallmarks that show age-dependent changes.

Genomic instability was found to have the strongest association with skeletal muscle aging, followed by stem cell exhaustion (**Figure 1D, E**). Importantly, only these two hallmarks showed significant age-dependent changes. Genomic instability, characterised by the accumulation of DNA damage and mutations, is a known cause of various functional declines associated with aging (Yousefzadeh *et al*., 2021). Our correlation network analysis among aging hallmarks (**Figure 1F, G**) further revealed that genomic instability was linked to most of the other hallmarks (**Figure 1H**).

### Splicing activity in aging is significantly associated with genomic instability

To investigate the influence of genomic instability on skeletal muscle aging, we thoroughly examined the pathways that interact with it. Therefore, we devised a method known as dGSEA (refer to the Methods section). This approach tests for pathway enrichment based on proteins (targets) that correlate with a specific factor or pathway of interest (source) (**Figure 2A**). Similar to traditional GSEA, the interpretation is that a positively enriched pathway indicates the function of proteins positively correlated with the ‘source’, while a negatively enriched pathway signifies the function of proteins that are negatively correlated. In this study, 15 proteins associated with genomic instability were designated as the ‘source’, and their correlated pathways were explored.

**Figure 2.**
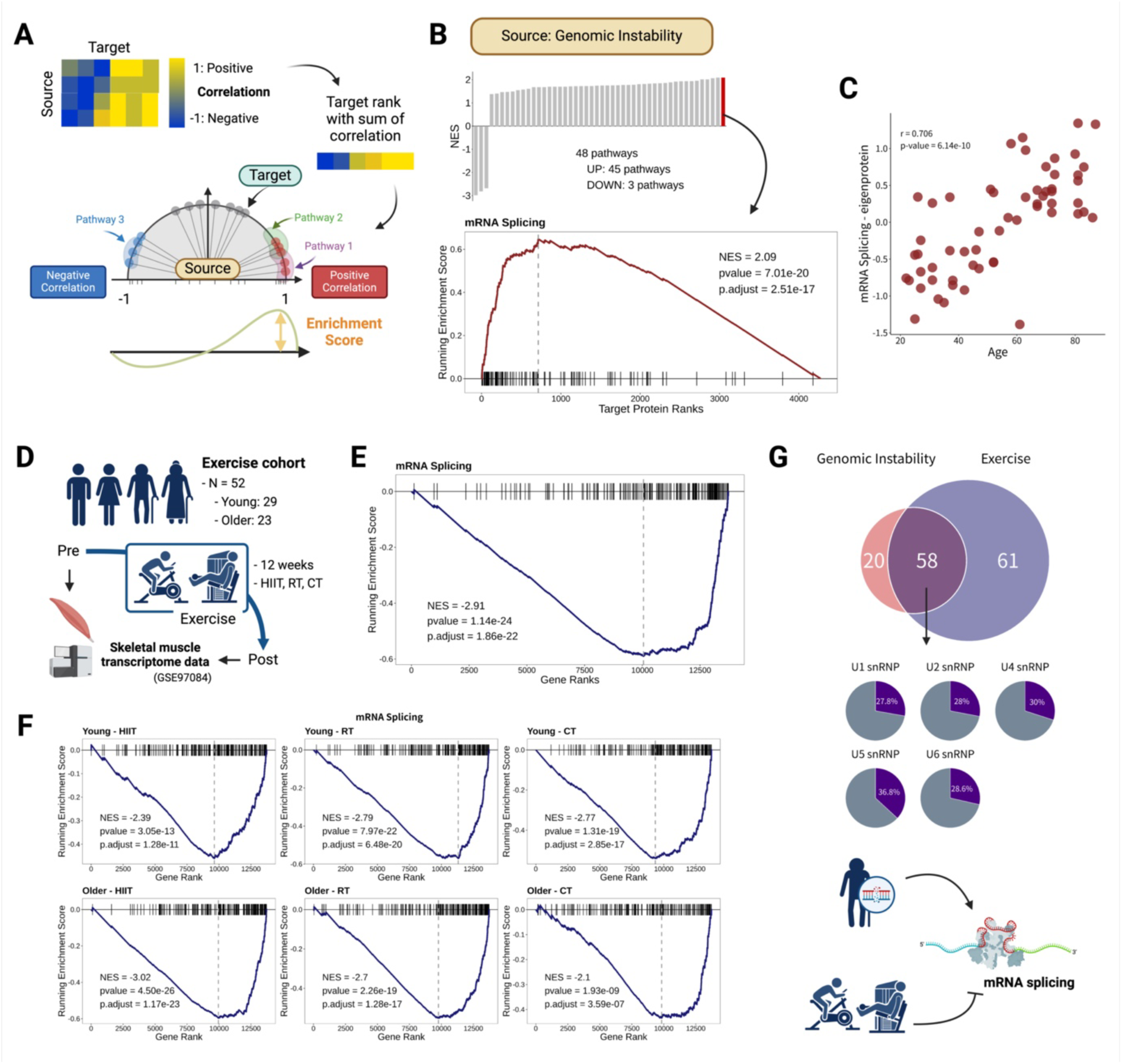
Contrasting effects of mRNA splicing regulated by genomic instability and exercise. **A)** Overview of the directional Gene Set Enrichment Analysis (dGSEA) method. This approach identifies pathways associated with specific ‘source’ molecules. The ‘target’ molecules associated with the ‘source’ are determined by computing the sum of their correlations, creating weights for each ‘target’ that are then used in the GSEA algorithm to identify involved pathways. The enrichment score indicates the level of enrichment of the molecules positively or negatively correlated with the ‘source’. **B)** Pathways significantly and most strongly enriched by dGSEA, using the genomic instability molecular signature as ‘source’. The top panel shows the 48 significantly enriched pathways, with the mRNA splicing pathway exhibiting the strongest enrichment (bottom panel). **C)** Age-related variation in mRNA splicing eigenprotein from the leading edge of dGSEA. Age is plotted on the x-axis, with the eigenprotein level of the mRNA splicing pathway on the y-axis. **D)** Overview of skeletal muscle transcriptome data collected pre- and post-exercise training used in the analysis. **E)** GSEA results highlighting mRNA splicing enrichment based on changes in gene expression pre- and post-exercise training. **F)** GSEA results for mRNA splicing based on gene expression changes pre- and post-exercise training across six subgroups. The top panels represent the results of the young group, and the bottom panels represent the results of the older group, with high-intensity interval training (HIIT), resistance training (RT), and combined training (CT) from left to right, respectively. **G)** A comparison of mRNA splicing-associated molecules upregulated due to genomic instability in an age-dependent manner and conversely downregulated due to exercise (top panel). The distribution of overlapping mRNA splicing-associated molecules across different spliceosome components is shown in the bottom panel. Parts of the figures were created with biorender.com.

We discovered 48 pathways significantly correlated with genomic instability (**Table 1**). Notably, mRNA splicing was identified as the most strongly and positively correlated with genomic instability (**Figure 2B**). mRNA splicing is crucial for proteomic diversity (Liu *et al*., 2017), and its dysregulation has been proposed as a new hallmark of aging (Schmauck-Medina *et al*., 2022). In our analysis, six of the top ten enriched pathways were related to splicing (**Table 1**). Furthermore, mRNA splicing activity displayed a significant correlation with age (**Figure 2C**). Consequently, our subsequent analysis focused on mRNA splicing.

**Table 1.**
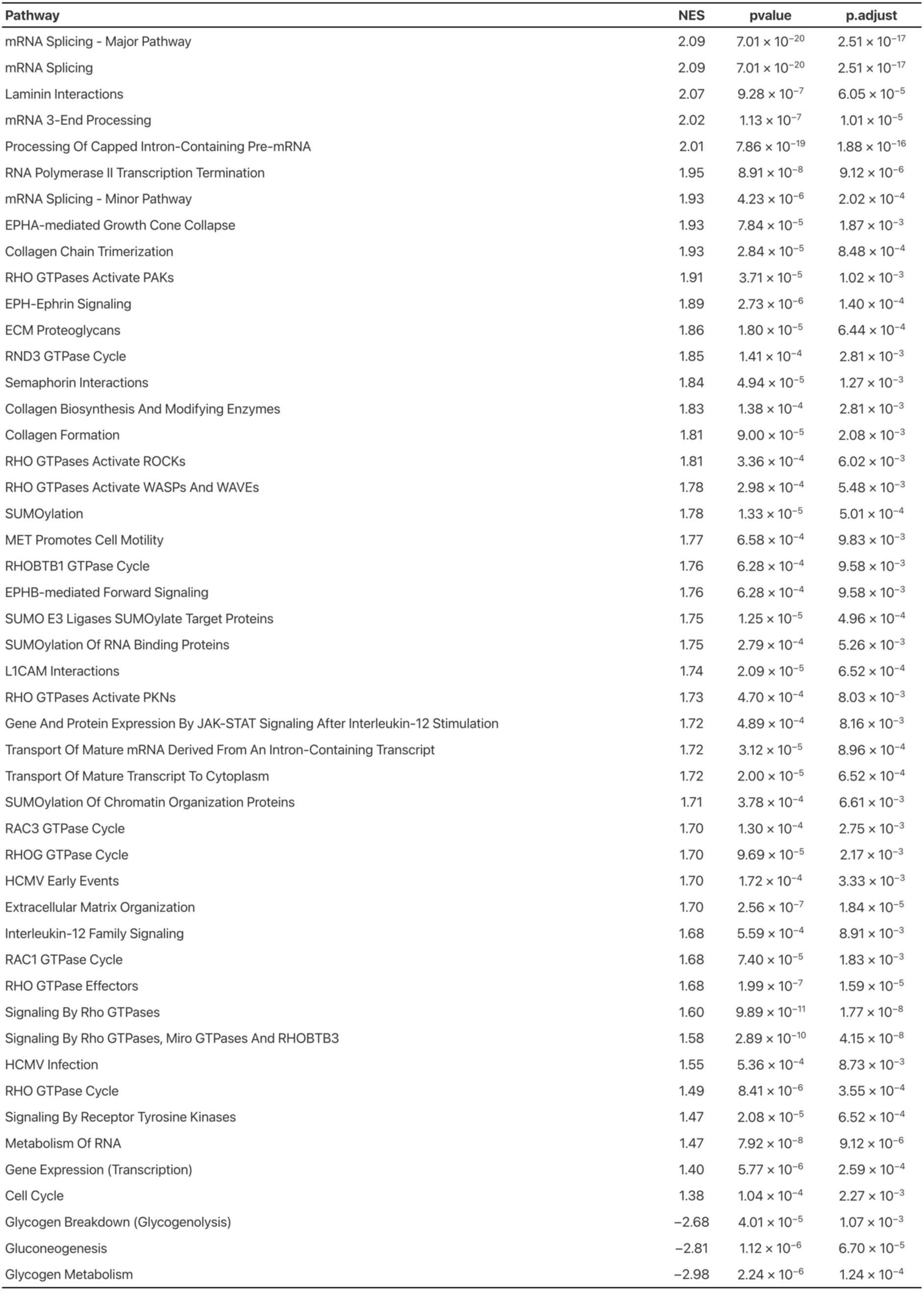
Pathways significantly correlated with genomic instability in skeletal muscle aging.

### Exercise suppresses the age-related increase in splicing activity

We then explored the impact of exercise on mRNA splicing activity. Human skeletal muscle RNA-seq data collected from healthy adults pre- and post-exercise training (Robinson *et al*., 2017) (GSE97084) (**Figure 2D**) were assessed. Gene signatures with altered expression pre- and post-exercise training were identified, and enrichment analysis was conducted using GSEA. Notably, exercise significantly downregulated mRNA splicing, in contrast to the aging signature (**Figure 2E**).

We further analysed whether the impact of exercise on mRNA splicing varied among the three exercise modalities (HIIT, RT, and CT) according to different age (young and older). The analysis revealed that exercise downregulated mRNA splicing activity across all exercise modalities and age groups (**Figure 2F**). There were 58 overlaps between proteins involved in mRNA splicing that were upregulated by genomic instability and genes downregulated by exercise, indicating a significant overlap with spliceosomes (**Figure 2G**).

### IR is the primary splicing event altered during the aging process

To assess the role of mRNA splicing in the anti-aging effects of exercise, splicing events, including SE, MX, A5, A3, AF, AL, and IR (see the Methods section), were identified using SUPPA2 (Trincado *et al*., 2018). The relative abundance of these splicing events is commonly denoted as PSI (Trincado *et al*., 2018) (**Figure 3A**). We estimated the PSI and analysed the age dependence of each splicing event through regression analysis, identifying IR as having the strongest correlation with age (**Figure 3B**). IR refers to a splicing event that retains introns that are typically removed by mRNA splicing, a process known to intensify with age and disease (Adusumalli *et al*., 2019; Mariotti *et al*., 2022). This observation was further validated using another RNA-seq dataset from an exercise cohort (Robinson *et al*., 2017) (GSE97084), which confirmed the age-related increase in IR (**Figure 3B**). These findings suggest that age-related changes in splicing regulation contribute to the rise in IR.

**Figure 3.**
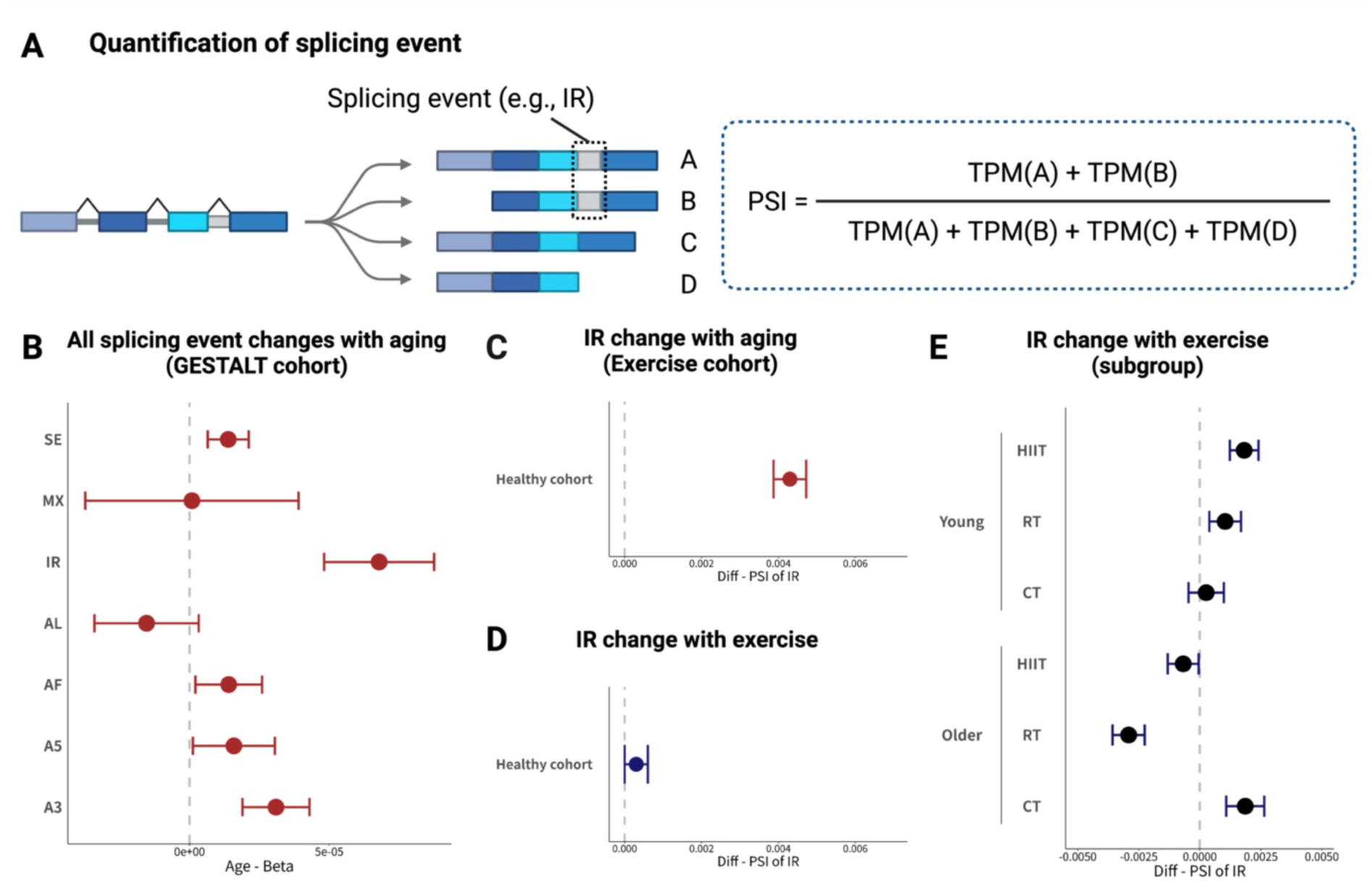
Aging and exercise have contrasting effects on IR. **A)** Overview of Percent Spliced In (PSI) calculation. PSI for each splicing event was estimated using TPM values of isoforms undergoing specific splicing events, as calculated using SUPPA2. An illustrative example demonstrates a gene with isoforms A, B, C, and D. **B)** Changes in all splicing events change with aging within the GESTALT cohort. The x-axis displays the regression coefficient of PSI when regressed on age, where a positive coefficient indicates a positive correlation of PSI with age, and vice versa. The vertical dashed line indicates no change. Circles denote means and error bars indicate standard errors. Abbreviations include skipping exon (SE), mutually exclusive exons (MX), alternative 5’/3’ splice sites (A5/A3), alternative first/last exons (AF/AL), and intron retention (IR). **C)** Changes in PSI for IR associated with aging, excluding exercise training, within the exercise cohort. The x-axis shows the difference in mean PSI of IR between the young and older groups, calculated as PSI (older group) - PSI (young group). **D)** Changes in PSI for IR pre- and post-exercise training within the exercise cohort. The x-axis reflects the difference in mean PSI of IR pre- and post-exercise training, calculated as PSI (post-exercise training, representing PSI (post-exercise training) - PSI (pre-exercise training). **E)** Subgroup analysis for changes in PSI for IR, considering age and exercise modality within the exercise cohort. Pre- and post-exercise training PSI changes for IR across three exercise modalities for both young and older groups are depicted. The x-axis shows the difference in mean PSI of IR pre- and post-exercise training, calculated as PSI (post-exercise training) - PSI (pre- exercise training). Parts of the figures were generated with biorender.com.

### Exercise modulates IR in an age- and modality-dependent manner

We further investigated whether exercise could mitigate aging effects on IR. Using the RNA-seq dataset from pre- and post-exercise training (Robinson *et al*., 2017) (GSE97084), we analysed the PSI values for IR. Contrary to the aging process, exercise induced minimal changes in IR levels (**Figure 3C**). However, the influence of exercise on IR varied based on age and the type of exercise performed. Our subgroup analysis revealed distinct IR responses to exercise among the six subgroups. Notably, RT exhibited the most potent inhibitory effect on IR within the older group (**Figure 3D**). HIIT also demonstrated a suppressive effect on IR in the older group, albeit to a lesser extent (**Figure 3D**). Conversely, in the young group, no exercise modality showed significant suppression of IR (**Figure 3D**). These results indicate that the ability of exercise to suppress IR is contingent upon both the age of the individuals and the type of exercise undertaken.

### Principal and alternative isoforms exhibit distinct patterns for IR during aging and exercise

We focused on the contribution of IR to biological processes by examining ‘functional isoforms’, which include principal and alternative isoforms (Rodriguez *et al*., 2022). Principal isoforms constitute the majority of the proteome and carry out the primary functions of each gene (Tress *et al*., 2017). We categorised IR occurring within principal isoforms as IR-P and within alternative isoforms as IR-A, according to the APPRIS database (Rodriguez *et al*., 2022) (**Figure 4A**).

**Figure 4.**
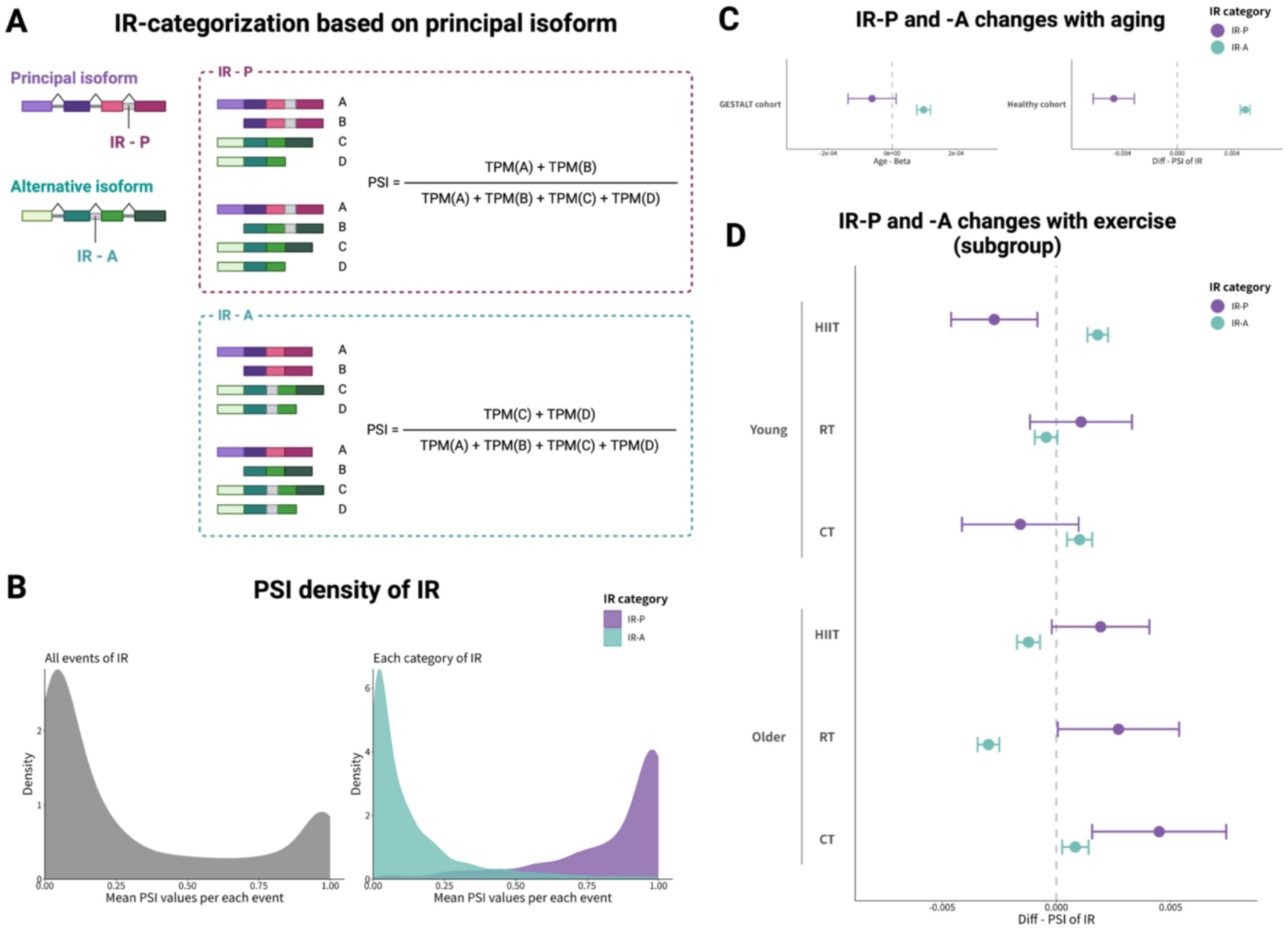
Directional changes in each IR category. **A)** Overview of IR categorisation and PSI calculation. Principal and alternative isoforms were classified according to the APPRIS database. IR occurring within principal isoforms was termed IR-P, and IR within alternative isoforms was termed IR-A. Examples include gene isoforms A, B, C, and D, with at least one of A or B being a principal isoform (top panel) and at least one being an alternative isoform (bottom panel). Approximate PSI calculations for principal isoform IR (IR-P) involve combining PSI values for various isoforms, including alternative ones, as PSI estimates are derived from TPMs based on reads containing IRs. **B)** PSI density distribution for IR. The right panel displays the PSI density for all splicing events, while the left panel focuses on the PSI density for each IR category, namely IR-P and IR-A. **C)** Age-associated changes in PSI for IR-P and IR-A within the GESTALT cohort (left panel) and pre-exercise training samples from the exercise cohort (right panel). The y-axis indicates age-related PSI changes, with the left column displaying the regression coefficient of PSI on age, and the right column showing the difference in PSI between the older and younger groups. Error bars denote the standard error for the age regression coefficient (left panel) or the PSI difference between groups (right panel). **D)** Exercise training-induced changes in PSI for IR-P and IR-A across six exercise cohort subgroups. Circles indicate means and error bars represent the standard error for either the age regression coefficient or the PSI difference between groups. Parts of the figures were generated with biorender.com.

To assess the impact of IR on the principal isoforms, we devised a post-hoc approximation method for calculating PSI (**Figure 4A**). Existing methods such as SUPPA2 calculate PSI based on read counts that include IR, with percentages derived from the TPM values, but they do not differentiate between isoform types, necessitating a post hoc approach for estimating the PSI of IR-P and IR-A. Our computational method aims to quantify the effect of IR on at least one principal isoform. If an IR, identified using SUPPA2 in a bulk isoform, includes at least one principal isoform, we treated its PSI as the PSI estimate for IR-P (**Figure 4A**). Conversely, IR-A was assessed when all isoforms with IRs were alternative isoforms (**Figure 4A**). The calculated PSI distributions for IR-P and IR-A revealed a clear bimodal distribution, highlighting the functional divergence between principal and alternative isoforms (**Figure 4B**).

Linear regression analysis of PSI for IR-P and IR-A using RNA-seq data from the GESTALT cohort (Ubaida-Mohien *et al*., 2019*b*) (GSE129643)) revealed a decrease in PSI for IR-P and an increase for IR-A with aging (**Figure 4C**). This pattern was also observed in the exercise cohort (Robinson *et al*., 2017) (GSE97084) when comparing the young and older groups, aligning with recent findings (Mariotti *et al*., 2022). We further explored the influence of exercise on IR-P and IR-A, analysing PSI changes in both pre- and post-exercise training across age groups (Robinson *et al*., 2017) (GSE97084). Notably, RT in the older group increased PSI for IR-P while decreasing it for IR-A (**Figure 4D**), effectively reversing the aging-associated pattern of IR alterations.

### Effect of IR on principal isoform expression

The influence of IR-P on changes in principal isoform expression was assessed using the correlation coefficient between age-dependent changes in principal isoform expression and PSI variations in IR-P. Additionally, we examined the correlation between exercise-induced changes in principal isoform expression and PSI in IR-P and IR-A. The analysis revealed that PSI changes in IR, due to aging and exercise, were significantly and positively correlated with alterations in the expression levels of the corresponding principal isoforms (**Figure 5A**). A significant negative correlation was observed for IR-A, in contrast to IR-P (**Figure 5A**).

**Figure 5.**
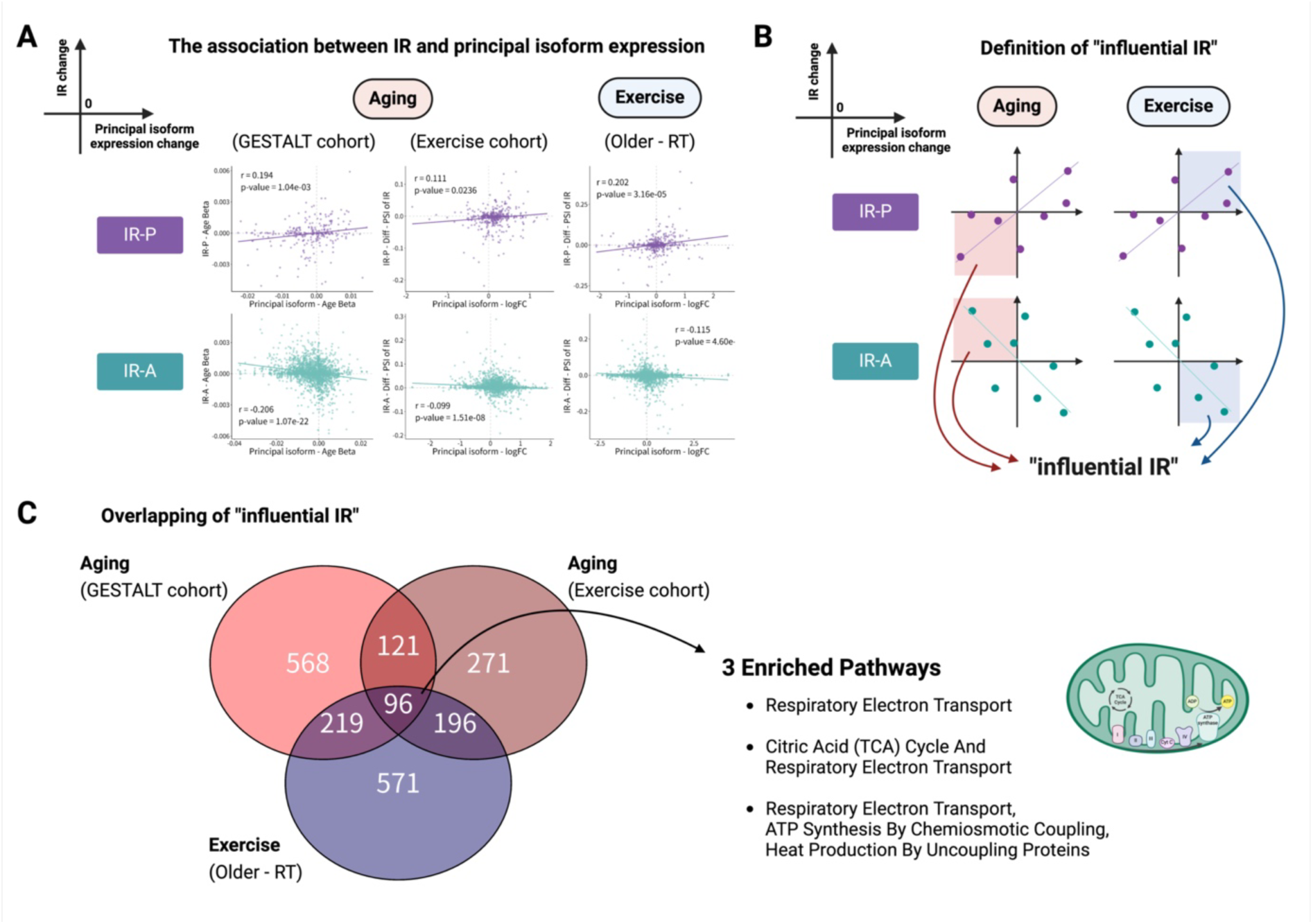
Mitochondrial function identified as an anti-aging process involving IR. **A)** Correlation between changes in IR-P or IR-A with aging and exercise training (Older - RT) and changes in principal isoform expression. Each point represents an isoform. The y-axis shows the PSI change in IR, with IR-P changes displayed in the upper panel and IR-A changes in the lower panel. The x-axis indicates changes in principal isoform expression. The left and middle panels show the changes in expression of principal isoforms with aging in the GESTALT and exercise cohorts, respectively, while the right panel depicts changes in principal isoform expression pre- and post-exercise training (RT) in the exercise cohort. Regression lines are plotted to illustrate trends. **B)** Computational identification of ‘influential IRs’ affecting principal isoform expression. Definitions derived from the scatterplot analysis in Figure 4A. For instance, ‘influential IRs’ associated with decreased principal isoform expression due to aging were identified in the lower left third quadrant (upper left panel), whereas ‘influential IRs’ contributing to improvements in principal isoform expression due to exercise training were identified in the upper right first quadrant (upper right panel). A similar methodology was applied for alternative isoforms, albeit with reversed correlation patterns (lower panels). **C)** A Venn diagram illustrates the overlap of ‘influential IRs’ between aging and exercise training (left panel), with the right panel highlighting enriched pathways among the overlapping IRs. Parts of the figures were generated with biorender.com.

### IRs involved in the anti-aging process are enriched in mitochondrial function

We investigated the biological processes associated with IR changes. Currently, no methods exist for predicting the biological functions that IRs may influence. We hypothesized that biological processes altering transcriptional output with aging, contrary to the effects of exercise training, could indicate the effects of exercise training on aging. Specifically, we suggested that the decreased expression of principal isoforms in older individuals might indicate functional decline, while the increased expression of these principal isoforms with exercise training might indicate functional recovery. Thus, we identified ‘influential IRs’ as those involved in such isoform expression changes between aging and exercise training (**Figure 5B**).

In both aging and exercise cohorts, we identified 1004, 684 “influential IRs” for aging and 1082 for resistance training, with 96 IRs common to both (**Figure 5C**). The 96 IRs that were common among them were used in the subsequent analysis (**Figure 5C**). Enrichment analysis of these IRs highlighted three mitochondrial function pathways, especially those related to the electron transfer chain (**Figure 5C**). Furthermore, genes enriched for mitochondrial function included NDUFS2, NDUFS3, NDUFA10, and NDUFV, all integral to mitochondrial complex I (**Figure 5C**).

## Discussion

In this study, we conducted a systematic analysis guided by aging markers, focusing on proteomic and transcriptomic data from young and older adults. Notably, mRNA splicing activity, linked to genomic instability, increases with aging in skeletal muscle but is mitigated by exercise. Additionally, a detailed examination of splicing events revealed that IR plays a key role in the anti-aging effects of exercise, with IR-associated principal isoforms being predominantly involved in the mitochondrial electron transport chain.

Genomic instability is recognised as a contributing factor to various functional declines observed with aging. Our findings affirm that genomic instability stands as the primary hallmark of skeletal muscle aging (**Figure 1C-E**). The origins of age-related increases in genomic instability are not entirely clear but are thought to be partly due to an accumulation of somatic mutations over time. Indeed, analyses have shown that somatic mutations in bulk skeletal muscle mRNA sequences and muscle satellite cells accumulate with age(Franco *et al*., 2018; Yizhak *et al*., 2019; García-Nieto *et al*., 2019). Following genomic instability, stem cell exhaustion was identified as the second most significant aging-dependent hallmark, corroborating these observations (**Figures 1C-E**).

Our study demonstrates that mRNA splicing activity, which increases with genomic instability during skeletal muscle aging, can be diminished through exercise training interventions. Recent discussions have implicated mRNA splicing dysregulation as a newly recognised hallmark of aging (Schmauck-Medina *et al*., 2022), and it is also considered a critical hallmark in muscular dystrophy, sharing many characteristics with skeletal muscle aging (Solovyeva *et al*., 2021). To explore the impact of age- and exercise-induced changes in mRNA splicing, we conducted a differential analysis of splicing events (**Figure 3A**). Aberrant splicing plays a role in various physiological and pathological processes (Bhadra *et al*., 2020). While event-level aberrant splicing has been documented in the context of aging skeletal muscles (Tumasian *et al*., 2021), reports on the effects of exercise on such splicing anomalies in skeletal muscles are limited. For our differential splicing event analysis, we utilised SUPPA2 (Trincado *et al*., 2018), which was highlighted for its efficacy in a recent comparative study of various methods (Mehmood *et al*., 2020). Our analysis indicates that IR changes with age more so than other splicing events (**Figure 3B**). Furthermore, exercise appears to reverse age-related changes in IR, suggesting that exercise may counteract aging through modifications in splicing mechanisms (**Figure 3C-E**).

From an anti-aging perspective, dietary restriction has been found to increase IR and contribute to longevity by coordinating nonsense-mediated decay (Tabrez *et al*., 2017; Rollins *et al*., 2019). Some research on longevity suggests that certain IRs directly contribute to the longevity phenotype (Huang *et al*., 2022). However, there is limited evidence on the role of IR in anti-aging interventions, and our analysis presents a possible IR-related anti-aging effect of exercise. Our findings also indicate that the IR response to exercise varies by age group and exercise modality **(Figure 3C-E**), with IR decreasing following HIIT and RT in the older group. Notably, IR reduction was particularly observed with RT in the older group, suggesting RT may counteract age-related IR changes. Conversely, our results demonstrate that IR increased with exercise in certain groups, such as the young HIIT group.

We also explored the functional implications of IR changes due to aging and exercise, focusing on the principal isoforms (**Figure 4C-D**). The functional roles of the principal isoforms, as defined in the APPRIS database based on protein structure, functional features, and conservation across species, are distinct from those of alternative isoforms (Rodriguez *et al*., 2022). A recent cross-tissue study on age-related IR changes suggested that IR exhibits contrasting changes depending on whether they involve principal or alternative isoforms (Mariotti *et al*., 2022). In our analysis, IR involving principal isoforms (IR-P) decreased with age, while IR involving alternative isoforms (IR-A) increased (**Figure 4C**), aligning with existing literature (Mariotti *et al*., 2022). Significantly, in the older group, RT appeared to reverse age-related changes in both IR-P and IR-A, differing from other exercise modalities (**Figure 4D**). Furthermore, our enrichment analysis of IR, whose changes impact principal isoform expression, identified mitochondrial function (**Figure 5C**), especially the electron transport chain, as a key area involved. This suggests a novel hypothesis that IR changes in mitochondrial function are involved in exercise-induced anti-aging mechanisms.

Finally, the limitations of our study warrant discussion. The hypotheses generated through *in silico* analysis rely on currently available public datasets, derived from two independent healthy human skeletal muscle cohort samples of relatively small sizes (N = 58 and N = 52) and limited diversity. Thus, these hypotheses require validation with more comprehensive cohort data on exercise interventions for aging. Furthermore, to establish causal relationships more robustly, experiments using mouse or other models are necessary. Moreover, this study has not confirmed whether the IR isoforms identified are present at the peptide level. Proteogenomic analysis (Nesvizhskii, 2014) could be instrumental in determining whether RNA-seq-detected splicing events correspond to peptides identified using mass spectrometry. Nonetheless, given the limited sample sizes for studies on human skeletal muscle aging and exercise, an *in silico* hypothesis-generation approach based on public data remains valuable for elucidating the relationship between aging and exercise.

## Additional information

### Data availability

All data used in this study are available from public repositories. Transcriptome data are available from GSE97084 and GSE129643, which are registered with Gene Expression Omnibus. Proteome data are available from PXD011967, which is registered in PRIDE.

### Author contributions

**HK:** Formal analysis, Investigation, Resources, Data Curation, Writing - Original draft. **HI**: Writing - Reviewing and Editing, Supervision. **YM:** Conceptualization, Methodology, Software, Investigation, Writing - Reviewing and Editing, Funding acquisition, Project administration, Supervision.

